# Predictive Modeling of Sleep Slow Oscillation Emergence on the electrode manifold: Toward Personalized Closed-Loop Brain Stimulation

**DOI:** 10.1101/2024.09.03.611113

**Authors:** Mahmoud Alipour, Sara C. Mednick, Paola Malerba

## Abstract

**Background:** Sleep slow oscillations (SOs), characteristic of NREM sleep, are causally tied to cognitive outcomes and the health-promoting homeostatic functions of sleep. Due to these known benefits, brain stimulation techniques aiming to enhance SOs are being developed, with great potential to contribute to clinical interventions, as they hold promise for improving sleep functions in populations with identified SO deficits (e.g., mild cognitive impairment). SO-targeting closed-loop stimulation protocols currently strive to identify SO occurrences in real time, a computationally intensive step that can lead to reduced precision (compared to post-hoc detection). These approaches are also often limited to focusing on only one electrode location, thus inherently precluding targeting of SOs that is informed by the overall organization of SOs in space-time. Prediction of SO emergence across the electrode manifold would establish an alternative to online detection, thus greatly advancing the development of personalized and flexible brain stimulation paradigms. This study presents a computational model that predicts SO occurrences at multiple locations across a night of sleep. In combination with our previous study on optimizing brain stimulation protocols using the spatiotemporal properties of SOs, this model contributes to increasing the accuracy of SO targeting in brain stimulation applications.

**Methods:** SOs were detected in a dataset of nighttime sleep of 22 subjects (9 females), acquired with polysomnography including 64 EEG channels. Modeling of SO occurrence was achieved for SOs in stage N3, or in a combination of stages N2 and N3 (N2&N3). We study SO emergence at progressively more refined time scales. First, the cumulative SO occurrences in successive sleep cycles were successfully fit with exponentials. Secondly, the SO timing in each individual was modeled with a renewal point process. Using an inverse Gaussian model, we estimated the probability density function of SO timing and its parameters μ (mean) and λ (shape, representing skewness) in successive cycles.

**Results:** We observed a declining trend in the SO count across sleep cycles, which we modeled using a power law relationship. The decay rate per cycle was 1.473 for N3 and 1.139 for N2&N3, with variances of the decay rates across participants being 1 and 0.53, respectively. This pattern mirrors the declining trend of slow wave activity (SWA) across sleep cycles, likely due to the inherent relationship between SWA and SO. Additionally, the SO timing model for N3 showed an increasing trend in the model parameters (μ, λ) across cycles. The increase rate per cycle followed a power law relationship with a rate of 0.83 and an exponential relationship with a rate of 4.59, respectively. The variances of the increase rates were 0.02 for μ and 0.44 for λ across participants.

**Conclusion:** This study establishes a predictive model for SO occurrence during NREM sleep, providing insights into its organization in successive cycles and at different EEG channels, which is relevant to development of personalized stimulation paradigms. These findings imply that personalized model parameters can be estimated by incorporating SO information in the first sleep cycle, and hence SO timing can be predicted before its occurrence with a probability distribution, enabling more precise targeting of SOs.

## 1. Introduction

Cortical slow oscillations (SOs, 0.5-1.5 Hz) have been experimentally linked to health and cognitive functions of sleep such as synaptic homeostasis [1, 2], glymphatic clearance [3], working memory improvements [4, 5], and consolidation of episodic memory [6-9]. Mechanistically, SOs are thought to contribute to sleep-mediated consolidation by supporting the coordinated reactivation of memory traces and hence mediating activity-dependent reorganization of selected synapses. In particular, while each SO comprises a hyperpolarized down-state associated with decreased cortical activity and a depolarized up-state characterized by increased cortical activity [10], the SO up state is considered crucial to these functional roles, as it supports coordination with other sleep oscillations involved in memory consolidation, such as sleep spindles and hippocampal ripples [11, 12].

Research has shown that episodic memory consolidation across the night benefits from SO enhancement with electrical or auditory stimulation [13-16], thus introducing the idea of sleep-based stimulation techniques as an intervention to enhance/support the functional outcomes of sleep. However, the strides in this approach haven’t consistently yielded success [17-20]. One potential explanation for this inconsistency lies in the stochastic nature of SO emergence on the scalp, in both timing and location, which limits their reliable detection. Beyond detection, precise targeting of a phase during SOs is also essential to effectiveness, with stimulation during up-state prolonging trains of SOs and/or amplifying SO amplitudes [15], while stimulation during down-state induces delay or termination of SO trains [21]. Therefore, the challenge lies not only in detecting SOs, but in doing so with enough anticipation that the stimulation could be accurately timed to coincide with the appropriate phase.

The experimental setup for closed-loop brain stimulation aimed at targeting brain oscillations necessitates several crucial components, including real-time access to ongoing brain activity, an accurate detection algorithm for brain oscillations, and a stimulation device. In the context of real-time stimulation, any delay introduced in the process—whether in signal acquisition, data transmission to servers, or the implementation of the detection algorithm—can significantly diminish the efficacy and precision of the stimulation. These delays, often spanning several hundreds of milliseconds, pose a considerable challenge. The detection process of SOs typically requires several hundred milliseconds [22], while the up-state of SOs lasts for only a few hundred milliseconds [23]. Consequently, inherent delay in the detection process, coupled with other sources of delay, limits precise targeting of SOs. Current techniques often rely on real-time SO detection, which may compromise the precision of stimulation delivery. To overcome this limitation, this study introduces a computational model that predicts SO occurrences across multiple EEG channels to improve the accuracy and efficacy of closed-loop brain stimulation techniques in targeting SOs during a night of sleep.

In this work, we introduce a novel modeling approach designed to predict SO occurrence and optimize the timing of brain stimulation interventions, contributing to improving their efficacy in targeting SOs during sleep to enhance cognitive function. Utilizing a dataset of sleep nights in 22 subjects, we leverage renewal point processes to model the timing of SO occurrence within and across cycles, building on personalized information derived from the first sleep cycle.

Hence, this study establishes a predictive model for SO occurrence during sleep, offering valuable insights for the development of personalized closed-loop brain stimulation paradigms.

## 2. Materials and Methods

### 2.1. The sleep EEG dataset

The present study utilizes a dataset of sleep polysomnography from the Sleep and Cognition Lab at University of California Irvine, led by Dr. Mednick. The dataset is introduced in detail in [24]. The dataset includes full-night sleep EEG data from 22 healthy volunteers (9 females) without psychological or neurological issues. EEG were recorded using a 64-channel cap placed according to the international 10-20 System at a 1,000 Hz sampling rate, which was subsequently down-sampled to 128 Hz. Out of the 64 channels, 58 recorded the electrical activity of the brain, while others served as reference, ground, and other biosignal channels (e.g., EOG, EMG). Sleep stages (Wake, N1, N2, N3, and REM sleep) were visually scored in 30-second epochs following the R&K manual [25], using the MATLAB toolbox HUME [26]. Participants all had a good quality sleep (basic sleep outcomes in Supplementary Table S1).

### 2.2. SO detection

To detect SO events, we employed an algorithm previously utilized in [24], closely following the criteria introduced by Massimini et al. [27] and Dang-Vu et al. [28] SO detection was performed at each electrode independently, excluding time intervals where artifacts ware identified. The algorithm relied on well-established conditions regarding the amplitude and duration of SO events, and a comprehensive description can be found in prior publications [24, 29]. EEG epochs containing two types of artifacts were excluded. First, 30-second epochs containing artifacts such as movements or arousals, as determined by the expert sleep scorer at the Sleep and Cognition Lab at the University of California, Irvine, were discarded. Second, epochs with muscle movement artifacts were algorithmically identified and excluded using two methods. Following Brunner and colleagues [30], we identified as artifacts 4-second-long time bins in which power in the 26.25-32 Hz range exceeded 4 times the median of the 45 surrounding bins (3 minutes around the epoch) Similarly, following Wang and colleagues [31], we identified as artifacts 5-second-long time bins in which power in the 4-50Hz range exceeded 6 times the median of all bins. The EEG referenced to contralateral mastoids was filtered within the 0.1–4 Hz range, and potential SOs were identified in artifact-free epochs of NREM sleep by analyzing each channel independently. Candidate SOs were segments between consecutive positive-to-negative and negative-to-positive transitions that also satisfied the following criteria: (1) the minimum wave amplitude was below or equal to 80 uV, (2) the voltage range between maximum and minimum values was at least 80 uV, (3) the time between the first and second zero crossing in the data fell within 300ms to 1000ms, and (4) the total duration of the candidate event was no more than 10 seconds. The pool of candidate SO events that met these parameters underwent further screening to eliminate potential artifacts. This involved computing the amplitude at the trough referenced to the average signal ±10 seconds around the minimum. Events at one electrode with an amplitude size 4 standard deviations above the mean of all events detected at that electrode were discarded. A secondary distribution of amplitudes, encompassing all events from all electrodes of a subject, was then created. Once again, events with amplitudes above 4 standard deviations from the mean were discarded. When selecting SOs occurring during a stage, only those with both the beginning and end falling within the sleep stage were considered.

### 2.3. Sleep cycle detection

For sleep cycle detection, we utilized the Feinberg and Floyd algorithm [32]. Briefly, this algorithm requires a minimum of 5 minutes for the REM period, except for the first cycle, and at least 15 minutes for the NREM period to prevent brief stage 2 epochs in the REM period from being considered as separate NREM periods. Additionally, the start of a cycle is determined by the start of stage 2. In our study, we consider only sleep cycles that include both REM and NREM periods. Therefore, we excluded the last sleep cycle of participants if they showed no REM period.

2.4. SO percentage measurement

This study investigates patterns in SO counts across successive cycles. To provide a comparable measure across participants, we use SO percentage, a normalized value for each participant. We measure the SO percentage during successive EEG epochs and sleep cycles using the following formulas:

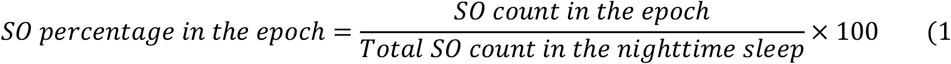

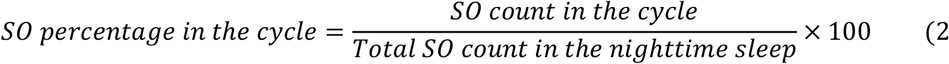

This study focuses on SO percentage during stage N3 and the combined stages N2 and N3 (N2&N3) of the NREM period. We measure SO percentage in 30-second epochs, which we encode as a time series of SO percentage in successive epochs. As we were interested in investigating the pattern of SO percentage across successive cycles and participants, we noted the variability in sleep stages count in sleep cycles and the resulting different lengths of NREM periods, which led us to the decision to scale the length of each NREM sleep, in order to achieve uniform treatability of the problem across all subjects. Supplementary Figure S1 displays the average epoch count across participants in the first four sleep cycles in our dataset. In each NREM period, we utilize a method outlined in previous work by one of the authors (Dr. Alipour) [33] to conduct a 100-point interpolation between each two successive SO percentages. This is followed by down sampling at a rate equivalent to the duration of the NREM period, resulting in a 100-point time series.

### 2.5. Modeling of inter-arrival time of SOs

This study models the timing of SO occurrence within and across cycles. First, we describe our modeling approach for the timing of SO occurrence within cycles. Then, by modeling the parameters of the model across cycles, we introduce a predictive modeling of SO occurrence using a point process [34]. Point process models can be used for event prediction by analyzing past occurrences of events and identifying patterns or trends that can help predict future events. They have broad applications in neuroscience for modeling neural activity [35]. The first step in predicting the occurrence of events is finding an appropriate probability distribution function (PDF) that can describe the probability distribution of event occurrence.

After determining the PDF for event occurrence, the inversion method, which is a commonly used technique for simulating random variates, can be employed to simulate event occurrences. The inversion method samples from a uniform distribution and then uses the inverse of the cumulative distribution function (CDF) to transform these uniform samples into samples from the desired distribution [36]. Since the CDF describes the probability that a random variable takes on a value less than or equal to a specified value [37], the inverse CDF describes the value of the random variable corresponding to a given probability. By generating random samples from a uniform distribution for the inverse CDF, we can simulate the occurrence of events [36].

This study employed an inverse Gaussian (IG) function as a PDF to model the probability distribution of SO occurrence inter-arrival times. IG has a broad application in modeling the probability distribution of neural behavior [38-40], and our preliminary analysis shows that IG outperforms other common functions for modeling the probability distribution of SO occurrence inter-arrival times in our dataset (Supplementary Table S2). Therefore, we selected IG to model the probability distribution. The IG model consists of two parameters: mean (μ) and lambda (λ), defined according to the following formula [41]:

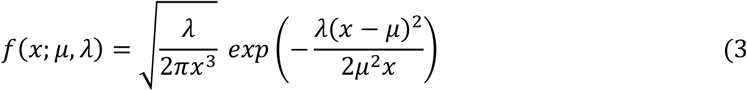

where *x* is the random variable. The optimal parameters of the IG model were determined by fitting an IG model to the distribution of the inter-arrival times of SOs within each sleep cycle. The CDF of the IG can be described by the following formula [42]:

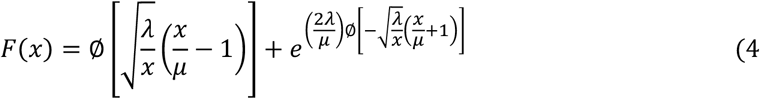

where ∅ is CDF of the standard normal distribution, as described by the formula below [43]:

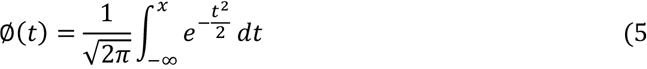

The CDF of normal distribution and its inverse are not available in closed form, requiring the use of numerical procedures [44]. Consequently, the inverse CDF of IG is also not available in a closed form. Therefore, we employed Matlab to estimate the inverse CDF of the IG distribution through numerical procedures, leveraging the error function [45] for estimating inverse CDF [46].

By fitting the IG model to the inter-arrival times of SOs for each sleep cycle, we obtain *μ* and *λ* parameters for each cycle. We repeat this process for all 56 EEG channels of 22 participants. Therefore, for each cycle, we obtain a 56 by 22 matrix for each of the *μ* and *λ* parameters. To develop a predictive model, our interest lies in identifying potential patterns in the variation of *μ* and *λ* parameters across successive cycles. By modeling *μ* and *λ* across cycles, we are able to predict the inter-arrival times of SO occurrence in later cycles based on the parameters estimated from the first sleep cycle. By averaging the mean value of *μ* and *λ* across participants and electrodes within each cycle, we obtain an average value 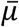 and 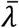 for each cycle.

Prior to averaging, sleep cycles with an SO count lower than a threshold for SO count of that cycle were excluded. This threshold was determined as the median of the lower quartile of SO counts across channels and subjects. Among the 56 EEG electrodes, eight electrodes including TP7, TP8, P1, P2, P5, P6, PO3, and PO4 were excluded from the modeling process due to limitations in SO count within cycles. For modeling the inter-arrival time of SO occurrence, we focused exclusively on N3.

To simulate the inter-arrival time of SOs, we generated random numbers from a uniform distribution. We then used the inverse CDF to convert those values to simulated inter-arrival times of SOs. Only inter-arrival times that exceed the refractory period for the given sleep cycle were accepted into the simulation. Similar to the refractory period in neuron firing rate [47], we defined a refractory period for our SO-generation process as the minimum time interval required for the resynchronization of neural activity to generate a new SO. We chose to use as refractory time the average of the minimum time delay between successive SO occurrences across all electrodes and participants. The simulation was built on an iterative process, including continued generation of simulated inter-arrival time values until the last inter-arrival time exceeded the cumulative summation of all inter-arrival times of SOs.

### 2.6. Model evaluation

We used three measures to quantify modeling error, including absolute difference error, Kolmogorov-Smirnov test (KS-test), and Kullback-Leibler divergence, employing bootstrapping 100 times. Absolute difference error is defined by averaging the difference between the predicted and real data times, divided by the real data time, as expressed in the following formula:

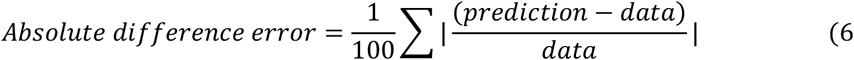

Kullback–Leibler divergence is a measure to quantify the dissimilarity between two probability distributions by explaining how one probability distribution diverges from another [48]. Here, we applied it to the comparison between the histogram of data and our simulation resulted from prediction. It can be defined using the following formula:

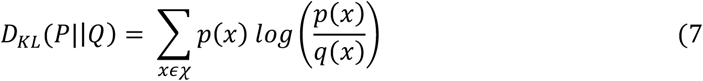

where P and Q are probability distributions on the same sample space *χ*, and x is the bin number in the histogram. p(x) and q(x) are probability density function of P and Q respectively. Values of this measure closer to zero indicate lower dissimilarity.

We used the KS-test to assess whether the distributions of our data and simulations could be derived from the same probability distribution. To obtain a cumulative value across multiple simulations, we defined a “distribution similarity” metric, as the fraction of 100 simulations which passed the KS-test (p-value > 0.05) when compared to a sample of real data. We interpret this measure as an indicator of similarity, ranging between 0 and 1 (after normalization), with values closer to 1 indicating higher similarity.

In addition to the three mentioned measures, we utilize a probability-probability (P-P) plot to visually assess the performance of the model by examining how closely a dataset aligns with a particular model. This method entails plotting the two cumulative distribution functions against each other. If they closely resemble each other, the data will appear as a nearly straight line [49]. Confidence intervals represent the lower and upper bounds of the range within which we expect the CDF of the data to lie with a certain level of confidence, 95% in this study. If the points in the P-P plot closely adhere to the diagonal line (y = x), it indicates a good fit between the distribution of the data and the model. Conversely, if a significant number of points in the P-P plot fall outside the confidence intervals, it may suggest a lack of fit between the distributions of the data and the model.

To represent the performance of the model in a P-P plot across all subjects, we followed the method outlined in reference [33] to standardize the CDF lengths of both the data and model to 250 points for each participant. Subsequently, we computed the mean CDF across all participants for both the data and the model. Plotting these mean CDFs against each other for each cycle produced the P-P plot. This resulting plot visually illustrates the average performance of the model across all participants in our study.

## 3. Results

We modeled SO occurrence across the night in two stages: first we identified the trend of SO percentage in successive cycles, then, we predicted SO occurrence within given cycles. In both steps, we defined a model that relied on information from the first sleep cycle. This approach allows us to estimate the parameters of the models in later sleep cycles by deriving them from measures acquired in the first sleep cycle.

### 3.1. SO percentage across cycles

We were firstly interested in estimating and modeling the cumulative trend of SO count decay across subsequent sleep cycles. For this analysis, we focused on the first four sleep cycles for each participant, since most participants had 4 or fewer cycles (Figure 1A) and cycles after the fourth had a very small amount of N3, which is the target of this modeling investigation. Consistent with known trends of slow wave activity (SWA), we observed a decrease in the SO percentage during successive sleep cycles, both in each participant (Figure 1B shows one example) and in a normalized averaged representation of cycle-by-cycle SO emergence across all participants (Figure 1C). When considering the cumulative amount of SOs found within each cycle, Figure 1D shows again the decreasing trend across them, which we modeled with a power law relationship:

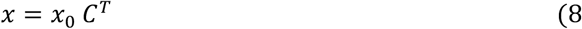

where *x* represents the SO percentage in a given cycle (a cumulative count across all cycle divided by the total count of SOs in the night), *x*_0_ is the SO percentage in the first cycle, C denotes the cycle number (1≤*C*≤), and T, the rate of decay, is equal to −1.473. The root mean squared error (RMSE) of the model equaled 0.0085. We modeled the across-cycle trend for each participant with a power law separately and found that the average decay rate across participants is −1.664 ±1.002. Thus, the SO percentage in cycles 2-4 can be estimated based on the SO percentage in the first cycle. By exploring the total SO percentage across sleep cycles during N2&N3, a similar decreasing trend is observed (Supplementary Figure S1). The decay rate for this trend is −1.139, and the RMSE of the model is 0.0064. The average decay rate across participants is −1.209 ± 0.726.

**Figure 1.**
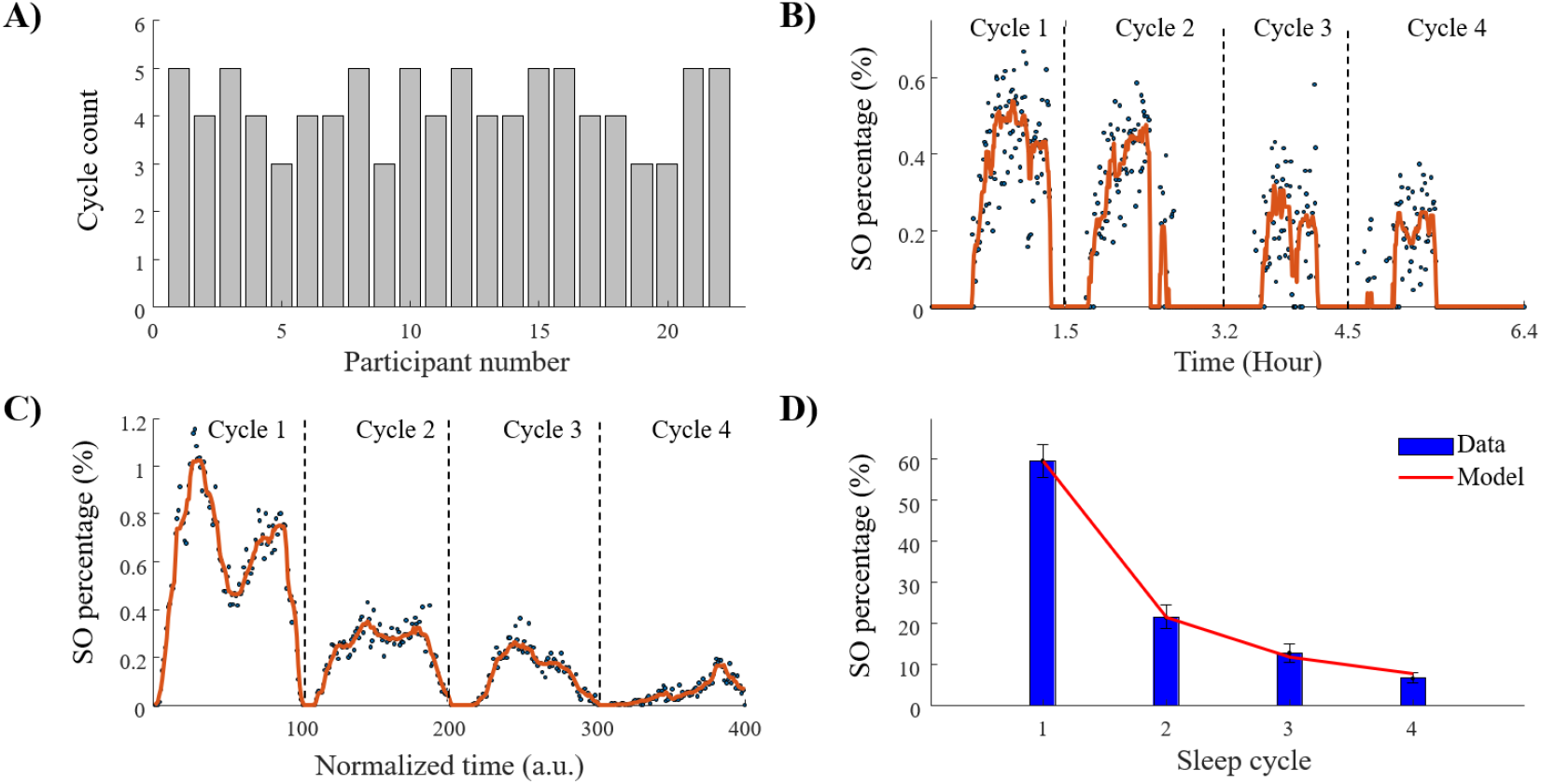
SO percentage across successive cycles. A) cycle count in different participants of our dataset. The x-axis shows participant number in the dataset. Most participants have 4 or fewer cycle. The average cycle count in our study is 4.2. B) SO percentage in successive epochs across cycles during N3 of the first participants in our dataset. The x-axis shows time per hour from sleep onset to end of fourth cycle. SO percentage in each epoch is depicted with dot, and the brown color represents the smoothed SO percentage obtained by averaging across 10 epochs using median. C) SO percentage across successive cycles during N3 by averaging across all participants. The x-axis shows normalized time, with cycle length equalized to 100 points. SO percentage is depicted by dots, and the brown color shows the smoothed SO percentage by averaging across 10 epochs. D) Total SO percentage during N3 across successive cycles obtained by averaging SO percentage across all participants. The x-axis shows sleep cycle, and error bars indicate standard error.

### 3.2. SO occurrence prediction

Our approach to predicting SO occurrences involved predicting the inter-arrival time of successive SOs and then estimating the timing of SO occurrence by cumulatively summing the inter-arrival times. Implementing the procedure outlined in section 2.5, for each cycle, we obtained IG parameters, including *μ* and *λ*, across 56 EEG channels of 22 participants, totally 56 × 22=1,232 pairs of parameters. Figure 2A schematically shows the matrix of *μ* corresponding to one sleep cycle. A different matrix was generated for *μ* and *λ* for each of the four cycles. Figures 2B and 2C show the distributions of *μ* and *λ* values for each matrix for the four sleep cycles, respectively. Figure 2D shows an increasing trend of the mean of all µ values in the corresponding matrix across successive cycles, which we modeled again with power law

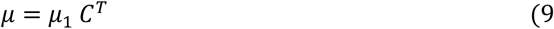

where *μ*_1_ represents the value of *μ* in first cycle, C is cycle number, and T, which is the increasing rate, equaled 0.8348. The RMSE of the model is 0.2381. By modeling μ for each participant separately, the average value of μ across participants is 0.831, with a variance of 0.44. Figure 2E shows the trends that emerge when the same procedure is applied to the distributions of λ values. We modeled the increasing trend for average λ with an exponential relationship as:

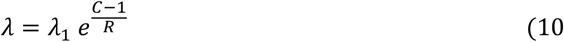

where *λ*_1_ is the value *λ* of in first cycle, C is cycle number, and R is increasing rate equal to 4.5893. The RMSE of the model equaled 0.0096. By modeling *λ* individually for each participant, the mean value of *λ* across all participants is 8.506, with a variance of 0.113. Relationships 9 and 10 demonstrate that the μ and λ values in subsequent cycles can be derived from the values of these parameters in the first sleep cycle.

**Figure 2.**
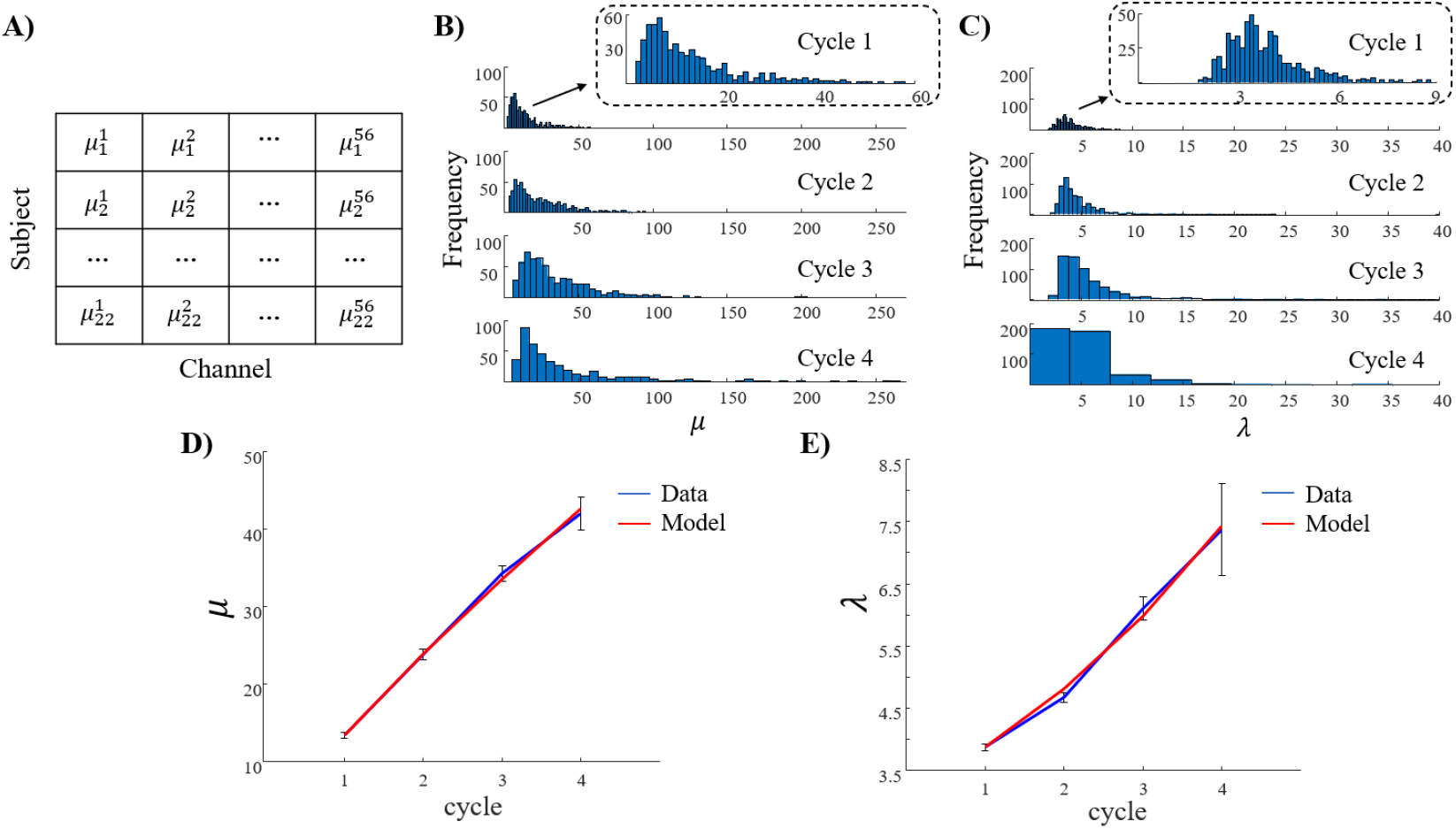
Model parameters of the IG as the PDF of inter-arrival time of SOs in successive sleep cycles. A) matrix of *μ* parameter for one sleep cycle across subjects and channels. Each cycle 1,232 has pairs of *μ* and *λ* values. B) Histogram of *μ* values for four sleep cycles. The x-axis shows *μ* value, and the y-axis shows the frequency or count of observations falling within each bin. C) Histogram of *λ*values for four sleep cycles. The x-axis shows *λ* value. In panel B and C, for clarification, a magnified picture is presented for first cycle. D) Average of *μ* values across subjects and channels in successive cycle. The increasing trend of *μ* across successive cycles can be model with a power law relationship. E) Average of values across subjects and channels in successive cycle. For averaging in panels D and E, the mean of 1,232 *μ* and *λ* values are calculated for each cycle. The increasing trend of λ across successive cycles can be model with an exponential relationship. Error bars in panels D and E represent the standard error.

To quantify model performance, we leveraged three measures: absolute difference error, distribution similarity, and Kullback-Leibler divergence (Table 1). To ensure robustness, each measure was subjected to 100 times bootstrapping. Evaluation was conducted based on the estimated parameters in cycles two to four, utilizing the parameters from the first cycle as a reference. Both absolute difference error and Kullback-Leibler divergence showed an increase across cycles, indicating that in simulations error increased with cycle number. Distribution similarity showed the maximum similarity between simulation and data in the third sleep cycle.

**Table 1.** Absolute difference error, distribution similarity and Kullback-Leibler divergence using 100 times bootstrapping.

|  | Cycle 2 | Cycle 3 | Cycle 4 |
| --- | --- | --- | --- |
| Absolute difference Error | $0.1774 \pm 0.0029$ | $0.1939 \pm 0.0030$ | $0.2941 \pm 0.0067$ |
| Distribution similarity (%) | $0.5079 \pm 0.0398$ | $0.7061 \pm 0.0435$ | $0.6343 \pm 0.0346$ |
| Kullback-Leibler divergence | $0.0092 \pm 0.0008$ | $0.0118 \pm 0.0013$ | $0.0124 \pm 0.0011$ |

To visually demonstrate the performance of the model, Figure 3 depicts a simulation of SO occurrence in third cycle of participant 4 in our dataset.

**Figure 3.**
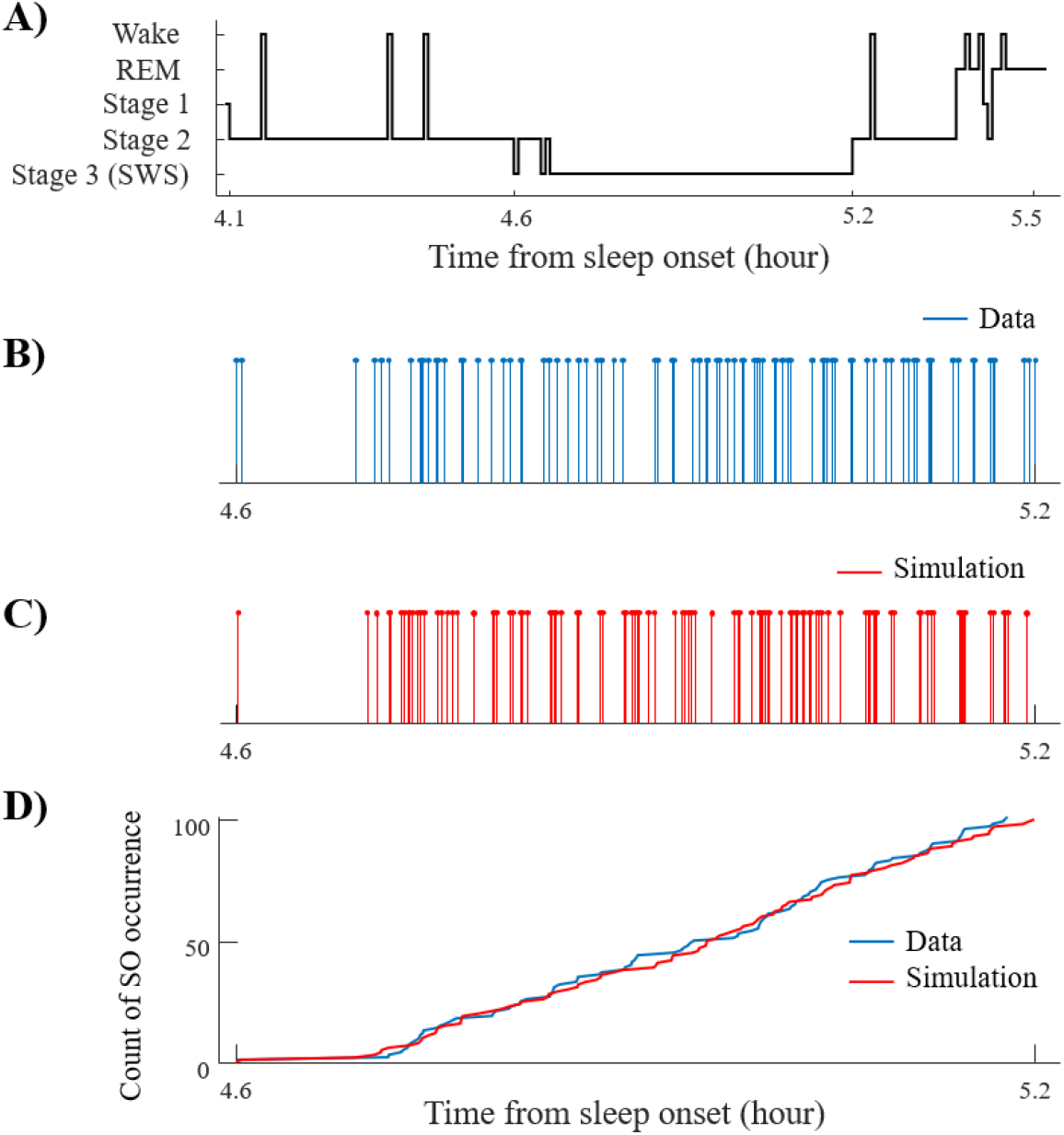
Simulation of SO occurrence in channel Fz during third cycle of one example participant in our dataset. A) Hypnogram of one participant from the start to the end of the third cycle. The x-axis represents time per hour from sleep onset. Note that EEG epochs with major body movements are shown as wake. B) SO occurrence in data. The count of SOs in this portion of data is 101. C) Simulation of SO occurrence with. *μ*=8.93 and *λ*=3.87. D) Timing of SO occurrence in data and one simulation of the third sleep cycle.

We also leveraged probability-probability (P-P) plots across successive sleep cycles, accompanied by a 95% confidence interval, to gain a sense of how close the simulations performed compared to data. Using the procedure outlined in section 2.6, within each cycle, the length of the CDF of both the data and the model was equalized to 250 points and then averaged across all participants. Closer proximity of points to the diagonal line indicates higher accuracy of the model in predicting SO occurrences. As illustrated in Fig 4, the majority of points fell within the confidence interval, demonstrating an acceptable goodness of fit between our model and the data.

**Figure 4.**
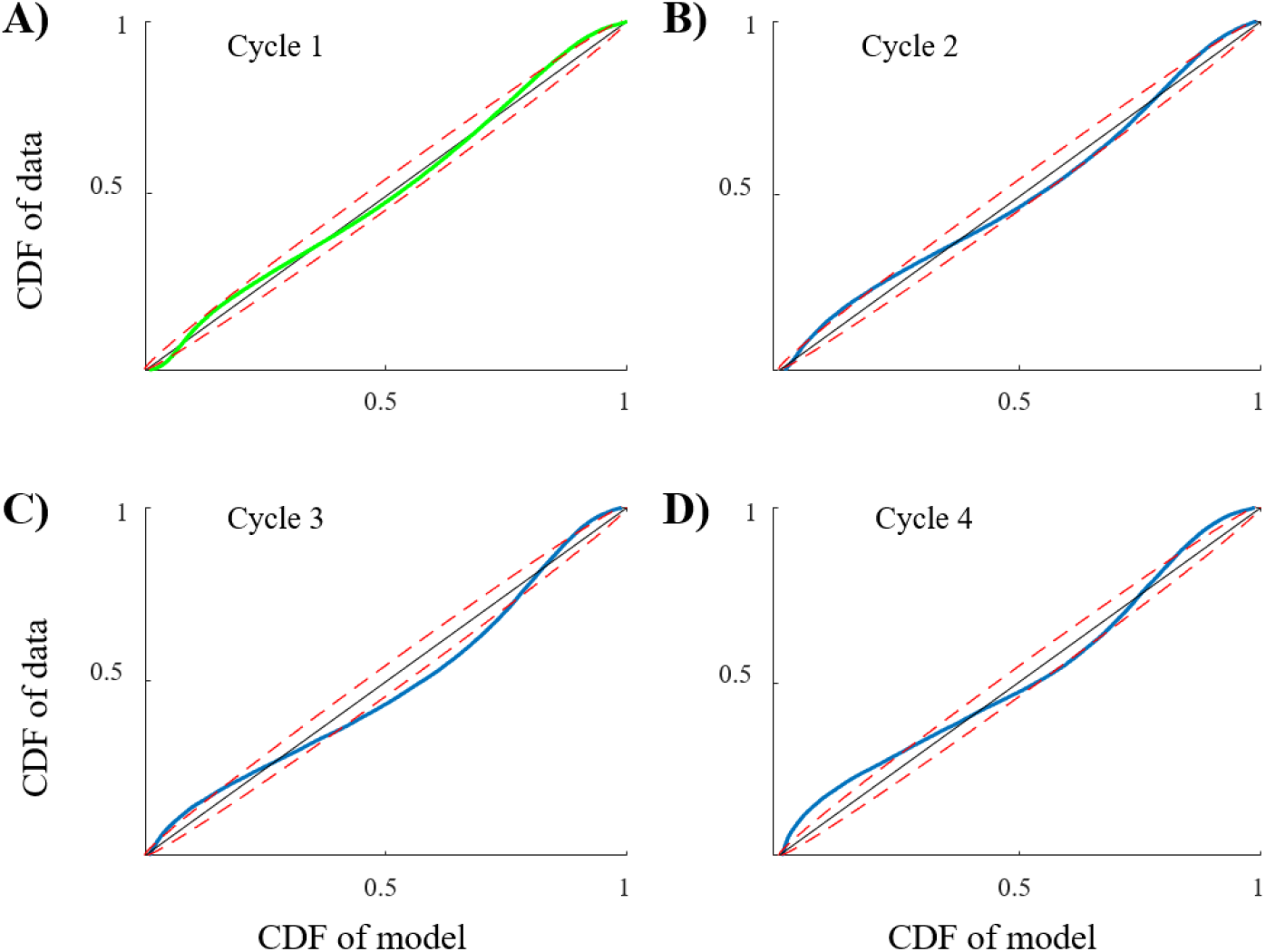
The P-P plot of data and simulation in successive sleep cycles. The y-axis shows CDF of data, while the x-axis represents CDF of the model. The red dashed line indicates 95% confidence interval. Panels A to D depict P-P plot for cycles 1 to 4, respectively. In Panel A, shown in green, the model parameters for the inter-arrival time of SOs in the first cycle are determined by fitting an IG model to the data. In subsequent cycles, shown in blue, the model parameters are estimated using relationships 9 and 10.

## 4. Discussion

This study presents a new approach to predicting when SOs will occur during sleep, using information from the first sleep cycle to guide predictions throughout the night. We introduce a modeling approach simulates SO emergence across the sleep night based on parameters derived from activity in the first sleep cycle. Our model captures how SOs gradually decrease in frequency (events per second) across sleep cycles as the night progresses, using a power law function dependent on the SO percentage in the first cycle. We model specific timing of SOs within each cycle by using point processes built on the distribution of SO inter-arrival times (represented with IG). Data showed that parameters μ and λ in the IG model changed across sleep cycles with an increasing trend, which we modeled with power law and exponential relationships, respectively. These findings shed light on the dynamics of SO occurrence across sleep cycles during nighttime sleep, which is important for developing more precise brain stimulation techniques that could enhance sleep quality and cognitive function.

Previous research has shown an exponential decreasing trend in the power of SWA across successive cycles [50], which serves as an index of sleep pressure (Process S) in sleep homeostasis and is well modeled in the literature [51-55]. Since fundamental dynamics of SOs contribute to the power of SWA, we expected to observe a reduced count of SOs in successive cycles. This study expands on the SWA-based knowledge to confirm substantially similar trends in SOs and to explicitly quantify the relationships between SO counts across subsequent sleep cycles.

Our model estimates the timing of SOs through by identifying the start of the transition from down state to up-state, which is a desirable target SO phase, given research has shown that stimulation targeting the SO up-state promote SO power and coordination with spindles [16, 56]. Results from this modeling study show that SO timing can be predicted (post-hoc) across the second to fourth sleep cycle by deriving parameters from the first sleep cycle. We suggest that this modeling strategy could be integrated in existing approaches that target SO occurrence in closed loop to enhance their performance. For example, adaptive weighting algorithms could be designed to integrate SO prediction and real-time SO detection. Furthermore, the potential of our model to estimate the timing of SOs using individualized parameters could contribute to approaches of personalized closed-loop brain stimulation during sleep. Such personalized approaches might be necessary to applications in clinical populations, or with respect to applications in older adults, given the temporal window for stimulus delivery has been found to shrink with aging [57]. Furthermore, given the pivotal role of SOs in cognition, there is potential in studying the trend of SOs across sleep cycles in cognitive disorders to explore any deviation from the observed trend in a normal cohort as a biomarker for the disorder. Our modeling study introduces a pair of parameters that can be estimated in each sleep cycle, which could provide objective quantitative measurements to subtle differences often discussed in the literature in atypical development [58, 59].

The current study had several limitations. Firstly, as our model is derived from a relatively small dataset, it represents an initial modeling step that requires replications in future studies with larger datasets. Secondly, our model does not represent activity past the fourth sleep cycle, despite some participants experiencing more than four cycles. The rationale behind this decision was the sparse occurrence of N3 after the fourth cycle in these participants. Future research could evaluate the cost/benefit of extending model fitting approaches beyond the fourth cycle, especially when considering modeling as contributing to the design of personalized stimulation paradigms. Another model limitation was our choice to derive a representative trend in model parameters as averaged across channels, assuming equal weighting for all channels during averaging. Since it is known that there is a higher frontal density of SOs [60], it is possible that a choice of modeling that allowed more impact to frontal channels compared to other channels, for example via weighted averaging, could yield improved results in future studies. Finally, our model makes some approximations on the relevance of time-scales that regulate SO emergence during the night. Due to the time dependency of sleep cycles, our model is time-dependent across cycles. However, within each cycle, our model is time-independent, meaning the rate of event generation doesn’t change over time. Additionally, while our model is memoryless, we attempted to mitigate this limitation by introducing a refractory period. As such, this model captures the fundamental dynamics of SO emergence in the second to fourth sleep cycle across the night, but it does not capture mathematically all the more subtle time and space dependent interactions that are the hallmark of an individual’s sleep EEG [61]. It is likely that more complex modeling would be required for that. Nonetheless, this crucial first step has identified some fundamental modes that more detailed models can build upon.

## Author contributions

### Mahmoud Alipour

Methodology, Data curation, Conceptualization, Formal Analysis, Software, Visualization, Writing – original draft. **Sara C. Mednick**: Writing – review & editing, Funding acquisition, Resources. **Paola Malerba**: Conceptualization, Funding acquisition, Investigation, Resources, Writing – review & editing, Methodology, Project administration, Supervision, Validation.

## Acknowledgements

NIH grant (R01 AG046646) to S.C.M. supported acquisition of data analyzed in this work.

