## Supplementary material for "Predictive Modeling of Sleep Slow Oscillation Emergence on the electrode manifold: Toward Personalized Closed-Loop Brain Stimulation"

**Tables**

Table S1. Sleep outcomes for our 22 participants. WASO: wake after sleep onset. For each value, time is reported in minutes.

| | |
|---|---|
| Total Sleep Time | $465.36 \pm 7.76$ |
| N1 | $22.1591 \pm 10.40$ |
| N2 | $199.75 \pm 25.92$ |
| N3 | $126.68 \pm 31.61$ |
| REM | $102.07 \pm 14.96$ |
| Sleep Onset | $14.04 \pm 6.71$ |
| WASO | $5.30 \pm 4.75$ |

Table S2. Performance of seven functions in modeling the inter-arrival times of SO occurrences in our dataset. We evaluated the performance of functions by calculating the ratio of p-values from the KS-test greater than 0.05 over 100 data simulations. For each data simulation within each cycle, we fitted the function to the data of that cycle. Values closer to 1 indicate a higher similarity between the data and the model. As can be seen, the inverse Gaussian function shows the highest value among the other functions.

| Function | Cycle 1 | Cycle 2 | Cycle 3 | Cycle 4 | Average across cycles |
|---|---|---|---|---|---|
| Exponential | 0.02 | 0.19 | 0.79 | 0.600 | 0.386 |
| Gamma | 0.10 | 0.53 | 0.93 | 0.860 | 0.618 |
| Inverse Gaussian | 0.31 | 0.71 | 0.90 | 0.880 | **0.708** |
| Logistic | 0.10 | 0.58 | 0.86 | 0.830 | 0.598 |
| Normal | 0.18 | 0.70 | 0.93 | 0.910 | 0.694 |
| Poisson | 0.03 | 0.55 | 0.90 | 0.870 | 0.592 |
| Weibull | 0.06 | 0.63 | 0.92 | 0.900 | 0.640 |

**Figures**

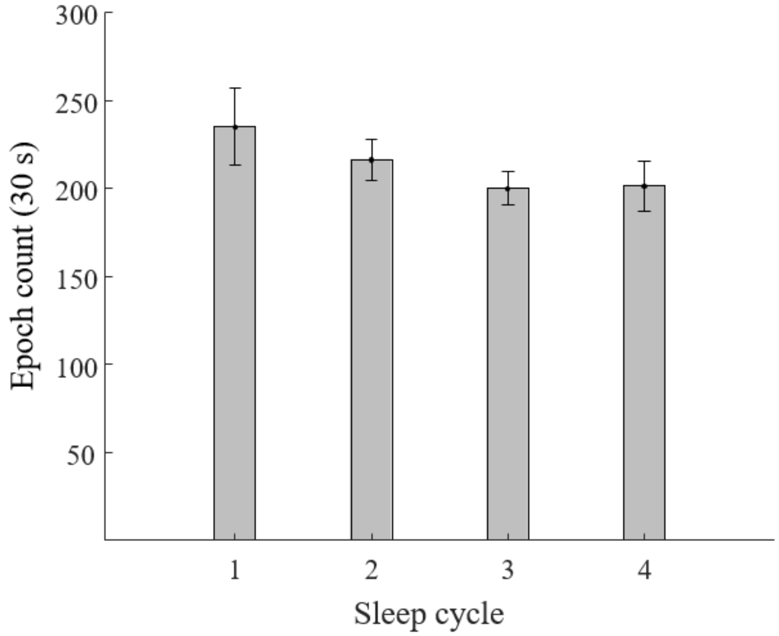


Figure S1. Average count of 30s EEG epochs across participants in sleep cycles. x-axis shows first four sleep cycles number and error bar shows standard error.

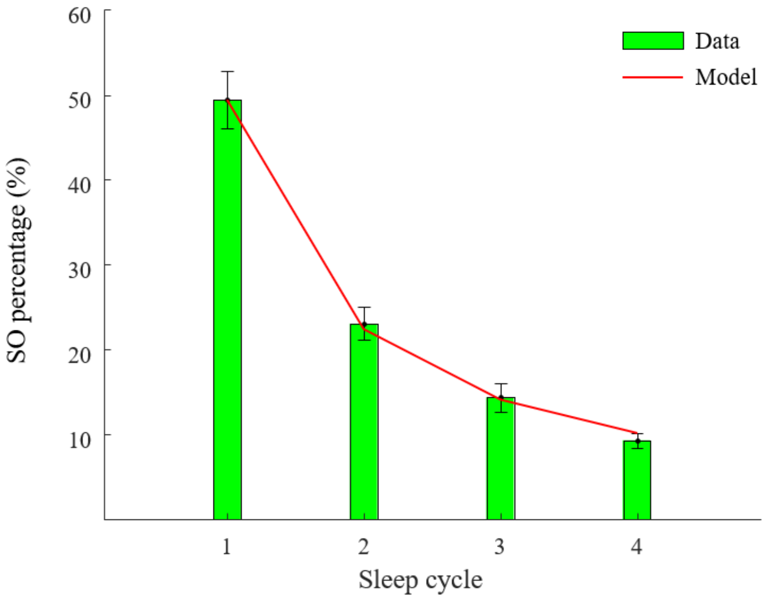


Figure S1. Total SO percentage during N2&N3 across successive cycles by averaging SO percentage across all participants. x-axis shows sleep cycle and error bar shows standard error. The decreasing trend in SO percentage across cycles can be described by a power law model with decreasing rate equal to -1.139. The RMSE of the model is 0.0064.
